# The X-Linked Intellectual Disability gene, *ZDHHC9*, is important for oligodendrocyte subtype determination and myelination

**DOI:** 10.1101/2023.08.08.552342

**Authors:** Rocio B. Hollman, Angela R. Wild, Timothy P. O’Leary, Andrew J. Thompson, Stephane Flibotte, Tashana O. Poblete, Angie Peng, Jason C. Rogalski, Gurmaan Gill, Shernaz X. Bamji

## Abstract

Two percent of patients with X-linked intellectual disability (XLID) exhibit loss-of-function mutations in the enzyme, ZDHHC9. One of the main anatomical deficits observed in these patients is a decrease in corpus callosum volume and a concurrent disruption in white matter integrity. In this study, we demonstrate that deletion of *Zdhhc9* in mice disrupts the balance of mature oligodendrocyte subtypes within the corpus callosum. While overall mature oligodendrocyte numbers are unchanged, there is a marked increase in MOL5/6 cells that are enriched in genes associated with cell adhesion and synapses, and a concomitant decrease in MOL2/3 cells that are enriched in genes associated with myelination. In line with this, we observed a decrease in the density of myelinated axons and disruptions in myelin compaction in the corpus callosum of *Zdhhc9* knockout mice. RNA sequencing and proteomic analysis further unveiled a reduction in genes and proteins essential for lipid metabolism, cholesterol synthesis, and myelin compaction. These findings reveal a previously under-appreciated and fundamental role for ZDHHC9 and protein palmitoylation in regulating oligodendrocyte subtype determination and myelinogenesis, offering mechanistic insights into the deficits observed in white matter volume in patients with mutations in *ZDHHC9*.

## Introduction

ZDHHC9 is one of a family of 23 palmitoyl acyl transferases that catalyze *S*-acylation, the reversible addition of acyl groups onto proteins through a labile thioester bond(1). In the brain, palmitoylation is the most common form of *S*-acylation and directly influences protein trafficking, function, and stability(1). Loss-of-function (LOF) mutations in the *ZDHHC9* gene (which is located on the X-chromosome) have been reported in ∼2 % of male patients diagnosed with X-linked intellectual disability (XLID)(2–9), with ∼75% of these patients exhibiting epileptic comorbidities(10). Deletion of *Zdhhc9* in mice (Gene nomenclature: “*ZDHHC9”* in humans and “*Zdhhc9”* in mice) results in cognitive deficits that mirror those observed in human patients(11).

Our lab has shown that ablating *Zdhhc9* can profoundly reduce dendritic branching, disrupt the formation of inhibitory synapses and increase seizure-like activity(12). Magnetic Resonance Imaging (MRI) on patients(4–6, 10) and mice(11) lacking ZDHHC9 demonstrate a striking decrease in the volume of the corpus callosum, a large white matter tract that connects the two hemispheres of the brain(13). In accordance with this, single-cell (scRNAseq) studies reveal that *Zdhhc9* is most highly expressed in the corpus callosum and in myelinating oligodendrocytes(14). Notably, *Zdhhc9* is the most highly expressed palmitoylating enzyme in oligodendrocytes at postnatal day 23(15)—the period of peak myelination of the corpus callosum(16). Despite the fact that ZDHHC9 is highly expressed in the corpus callosum and that the volume of the corpus callosum is significantly attenuated in patients and mice lacking ZDHHC9, the role of ZDHHC9 in white matter development remains unexplored.

White matter development is dependent on the proper differentiation of oligodendrocytes, which comprise >70% of cells in the corpus callosum(17). Oligodendrocyte differentiation is a stepwise process that starts with oligodendrocyte precursor cells (OPCs), that differentiate into newly formed oligodendrocytes (NFOLs), then myelin forming oligodendrocytes (MFOLs), and finally mature oligodendrocytes (MOLs)(15). Recent scRNAseq studies have demonstrated further transcriptomic heterogeneity within the four main oligodendrocyte groups including 2 NFOL subtypes (NFOL1,2), 2 MFOL subtypes (MFOL1,2) and 6 MOL subtypes (MOL1-6)(15, 18) that can be distinguished by their various gene expression signatures. For example, MOL2/3 cells are enriched in myelin-related and cholesterol biosynthesis genes while MOL5/6 cells exhibit high expression of synapse and cell-cell adhesion genes(15).

Here, we show that although ablation of ZDHHC9 does not impact the overall number of oligodendrocyte lineage cells, it is required for the proper determination of MOL subtype proportions in the corpus callosum, with *Zdhhc9* knockout (KO) mice exhibiting a decrease in MOL2/3 subtype cells and a concomitant increase in MOL5/6 cells. The decrease in MOL2/3 cells that are transcriptionally enriched for genes associated with myelination was accompanied by a decrease in the number of myelinated axons and disruptions in myelin compaction and integrity in the corpus callosum of KO mice. In addition, we observed reduced expression of genes and proteins essential for lipid metabolism, cholesterol synthesis, and myelin compaction. Together, this study provides a mechanistic understanding of the role of ZDHHC9 in white matter and suggests disruptions in interhemispheric cortical communication in patients with XLID and loss of function mutations in *ZDHHC9*.

## Results

### Uniform reduction in corpus callosum width throughout the anterior-posterior axis in *Zdhhc9*-KO mice

Axonal organization in the corpus callosum is heterogenous and organized in a spatially segregated manner according to projection neuron origin and corresponding targets in the contralateral hemisphere(19) (Figure S1a). Despite this heterogeneity, we observed a robust and relatively homogenous reduction in the width of the corpus callosum, hippocampal commissure and external capsule in postnatal day 60 (P60) *Zdhhc9*-KO male mice at all regions analyzed along the antero-posterior axis (Figure S1a-f), as well as a relatively homogenous expression of *Zdhhc9* along the anterior-posterior axis (Figure S1g,h). To date, only male patients have been identified to have XLID with *ZDHHC9* loss of function mutations(2, 13, 20), and accordingly, only male mice were used in this study.

## Altered corpus callosum gene expression in *Zdhhc9*-KO mice

To investigate whether there are any molecular alterations associated with changes in callosal volume, we performed bulk RNAseq on the microdissected corpus callosum of control and *Zdhhc9*-KO mice (Figure 1a). We identified 173 differentially expressed genes (Dataset S1 for RNAseq raw data; Dataset S2 for the list of 173 differentially expressed genes organized by gene ontology {GO annotation}). Of the differentially expressed genes, 52 were upregulated and 121 genes were downregulated in *Zdhhc9*-KO mice (Figure 1b). The top 5 downregulated genes other than *Zdhhc9* included: *Slco3a1*, *Apln*, *Slc45a3*, and *Cyp51*. *Slc45a3* and *Cyp51* are involved in lipid metabolism(21, 22), and similarly, other genes linked to this process such as *Ldlr, Hmgcs1, Insig1, Sqle,* and *Msmo1* were also downregulated. Further, *Apln* has been shown to enhance oligodendrocyte differentiation(23). The top 5 upregulated genes included: *Svep1*, *Kit*, *Syt2*, *Fbxw15*, *Oprk1*. Interestingly, both *Kit* and *Oprk1* have been associated with promoting oligodendrocyte proliferation(24–26). Aside from *Zdhhc9* itself, there was no significant change in the expression of other ZDHHC enzymes or palmitoyl thioesterases (enzymes that catalyze the removal of palmitate), suggesting no compensatory changes in other enzymes that regulate palmitoylation at the transcriptional level (Dataset S1). Notably, of all the ZDHHC enzymes, *Zdhhc9* was the most highly expressed ZDHHC enzyme in the corpus callosum of control P60 male mice, in line with previous studies(14, 27) (Dataset S1).

**Figure 1.**
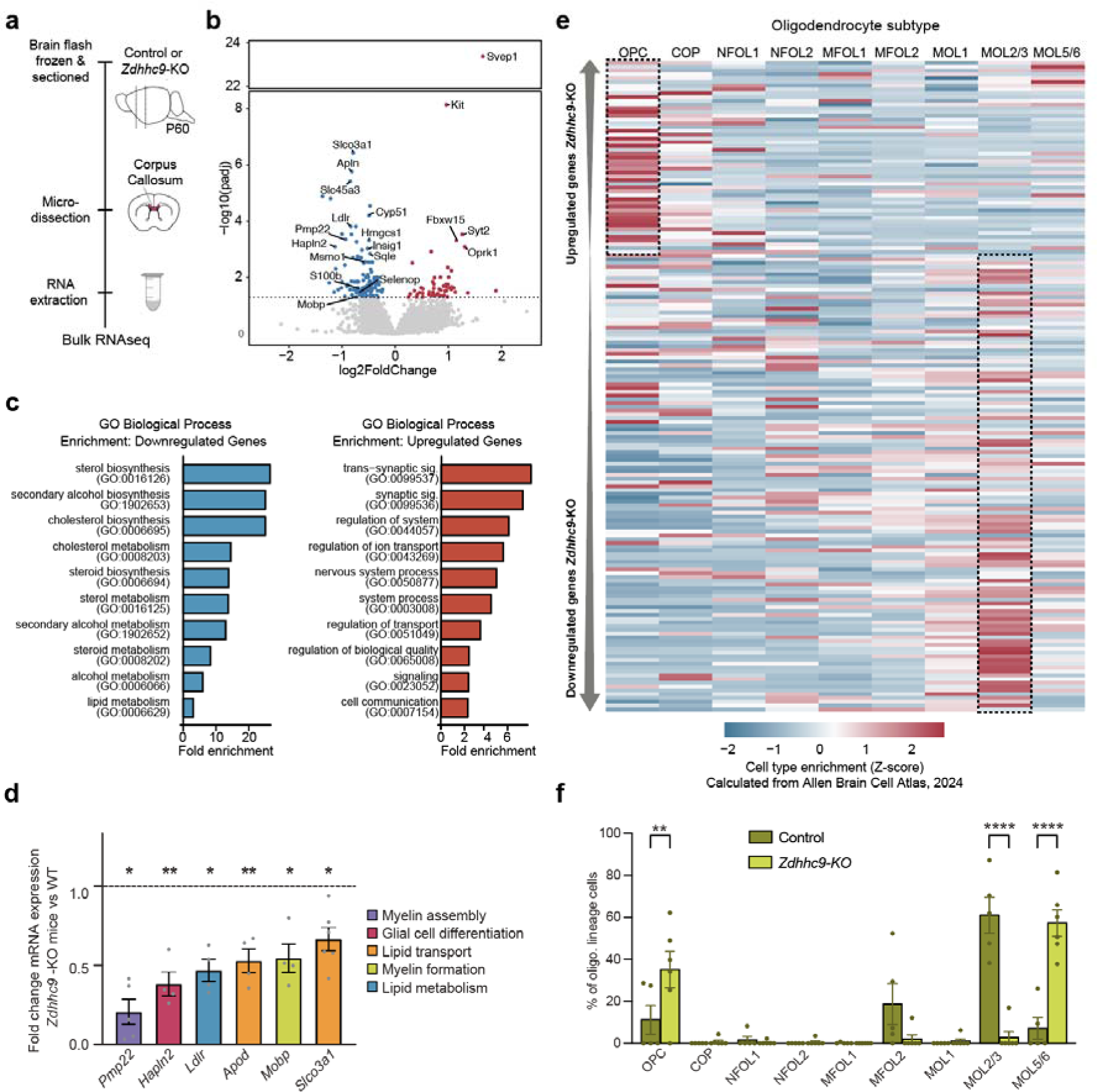
There is a change in mature oligodendrocyte gene expression and genes involved in lipid metabolism in the corpus callosum of *Zdhhc9*-KO mice. **a** Illustration showing pipeline for RNAseq analysis in P60 mice. **b** Quantification of fold change in gene expression (*n* = 5 *Zdhhc9-*KO mice, *n* = 6 control mice). Each point represents an individual gene; blue denotes genes that are downregulated, and red denotes genes that are upregulated in *Zdhhc9*-KO mice relative to controls. The horizontal line denotes the statistical significance threshold required for two tailed t-tests; p-values were adjusted to account for multiple comparisons. Top 5 downregulated and upregulated genes are annotated, along with genes of interest. **c** Enrichment analysis for Biological Process GO terms for both up- and downregulated genes. The ten GO terms with largest fold enrichment based on FDR value are shown. **d** Graph of RT-qPCR data for a subset of differentially expressed genes with known roles in myelination (*n* = 4-6 *Zdhhc9*-KO mice, *n*=4-6 control mice). Normalized expression of each gene relative to control (dotted line). **e** Heatmap showing the expression of the 173 differentially expressed genes within different oligodendrocyte subtypes using scRNAseq data from Allen Brain Cell (ABC) atlas. Genes are listed (top to bottom) by decreasing log_2_ Fold Change (i.e. from most to least upregulated and least to most downregulated). Dotted black boxes highlight predominant cell type enrichment of upregulated and downregulated genes. **f** Bisque bioinformatic estimation of oligodendrocyte cell type proportions in the corpus callosum, performed on bulk RNAseq data from this study and using Allen brain ABC atlas as a reference scRNAseq dataset.

To determine potential mechanisms and pathways that are perturbed by *Zdhhc9* ablation we completed PANTHER gene ontology (GO) enrichment analysis for biological processes on the differentially expressed genes. An analysis of downregulated genes revealed enrichment in numerous processes related to cholesterol and sterol metabolism and biosynthesis, whereas upregulated genes were enriched in pathways with the broad designation “signaling” (Figure 1c). Downregulated genes had a larger fold change compared to upregulated genes and smaller FDR values, indicating stronger enrichments in biological processes for downregulated genes.

We next examined the relationships between our 173 genes of interest using STRING (Search Tool for the Retrieval of Interacting Genes) protein-protein interaction network analysis (Figure S2). We identified two main clusters of largely downregulated genes with functions related to the immune system (highlighted in green) and cholesterol metabolism (highlighted in orange). The immune cluster included genes associated with microglia (such as *Ctss*, *C1qa*, *Ly86*, *Laptm5*, *Csf1r*; Figure 1b and Figure S2). Most genes in these clusters were downregulated, suggesting a potential impairment in both immune regulation and cholesterol metabolism in *Zdhhc9*-KO mice.

We next performed quantitative reverse transcriptase polymerase chain reaction (qRT-PCR) on six selected genes that were identified in key signaling networks in our RNAseq analysis, including myelin formation (*Mobp*), myelin assembly (*Pmp22*), glial cell differentiation (*Hapln2*), lipid transport (*Apod*, *Slco3a1*) and lipid metabolism (*Ldlr*). We confirmed decreased expression of all six of these genes in line with our RNAseq data, validating the robustness of our dataset (Figure 1d).

Changes in gene expression observed in bulk RNAseq data can be attributed to a variety of factors, including differential gene transcription within cells and variations in cell type proportions. We noted that several differentially expressed genes in *Zdhhc9*-KO mice were marker genes for mature oligodendrocytes (*Hapln2*, *Apod*, *Trf)*(15), indicating potential changes in oligodendrocyte cell type proportions. We therefore proceeded to investigate whether the genes exhibiting differential expression in the corpus callosum of *Zdhhc9*-KO mice were expressed in distinct oligodendrocyte subtypes. This was done by analyzing cell-type expression patterns using the latest comprehensive scRNA sequencing study of the adult (P60) mouse brain, conducted by the Allen Institute(28). Gene expression Z-scores were plotted for each oligodendrocyte subtype, with a high Z-score indicating enrichment within a particular cell type. We found that most genes that were up-regulated in *Zdhhc9*-KO mice were enriched in OPCs, while most down-regulated genes were enriched in MOL2/3 cells (Figure 1e). These results indicate that changes in gene expression might be in part due to changes in the subtype proportions of oligodendrocyte lineage cells.

To further investigate potential changes in oligodendrocyte cell type proportions, we used a bioinformatic bulk-RNAseq deconvolution method, Bisque, to estimate the cell type proportions in the corpus collosum of control and *Zdhhc9*-KO mice. Bisque analysis utilizes scRNA-seq data to generate a reference gene expression profile that can be applied to approximate cell subtype proportions within a bulk RNAseq sample(29). We utilized the scRNAseq study of the adult (P60) mouse brain from the Allen Institute as a reference dataset for Bisque. This analysis revealed an increase in OPCs and a decrease in MOL2/3 cells in *Zdhhc9*-KO mice (Figure 1f), mirroring our observation that upregulated genes are enriched in OPCs and down regulated genes are enriched in MOL2/3 cells (Figure 1e). Interestingly, we also observed an increase in the MOL5/6 subtype in *Zdhhc9*-KO mice (Figure 1f). This estimation of MOL5/6 may be influenced by the highly significant increased expression of *Svep1* (Figure 1b), which is a highly enriched gene in MOL6(15). These results point to potential changes in oligodendrocyte subtype proportions in the corpus callosum.

We also performed RNAseq on two additional micro dissected brain regions, including the primary visual cortex, in which seizure activity is present in *Zdhhc9* KO mice(12), and the optic nerve (Figure S3a,b), another large white matter tract. Although we did not identify any differentially expressed genes in the visual cortex, we identified 21 significantly downregulated and 14 significantly upregulated genes in the optic nerve of *Zdhhc9*-KO animals (Figure S3b; Dataset S3). Analysis of gene expression across oligodendrocyte subtypes revealed an enrichment of downregulated genes in the MOL2/3 subtype, as observed in the corpus callosum (Figure S3c; Dataset S3). Notably, five genes were significantly downregulated in both corpus callosum and optic nerve (*Apln*, *Selenop*, *Slc45a3*, *Pmp22 and S100b)* (Figure S3d). Among these, four genes (*Apln*, *Selenop*, *Slc45a3* and *Pmp22*) are most highly expressed in the MOL2/3 cell type and are known to be involved in oligodendrocyte differentiation and/or myelination(21, 23, 30, 31). Bisque approximation of cell type proportions again revealed a significant decrease in the MOL2/3 subtype of oligodendrocytes in optic nerve, indicating a decrease in this particular cell type that is common to both white matter tracts (Figure S3e). The shared alterations in gene expression observed in both the corpus callosum and optic nerve suggest broader changes in white matter gene expression due to *Zdhhc9* loss, in addition to more specific transcriptional changes in individual regions.

### Altered oligodendrocyte subtype proportions in *Zdhhc9*-KO mice

The changes in gene expression identified with RNAseq predict alterations in proportion of oligodendrocyte lineage cells in *Zdhhc9*-KO mice (Figure 1d). We therefore designed fluorescent *in situ* hybridization (FISH RNAscope) probes for the dominant MOL subtypes in the corpus callosum, guided by expression data from previously published scRNAseq and FISH studies(15, 28, 32)(Figure S4a). We chose marker genes with highly enriched expression in specific oligodendrocyte lineage cells, including *Hapln2* (MOL2/3 marker), *Ptgds* (MOL5/6 marker), and *C030029H02Rik* (MOL5/6 marker, Figure S4a). Importantly, within each cell type identified we found no difference in the mean probe coverage of each marker gene in control vs *Zdhhc9*-KO mice, confirming that these marker genes can be used to infer changes in cell number (Figure S4b). In line with our Bisque deconvolution analysis, there was a decrease in the proportion of MOL2/3 cells, with a concomitant increase in the proportion of MOL5/6 cells in the corpus callosum of *Zdhhc9-*KO mice (Figure 2a-c).

**Figure 2.**
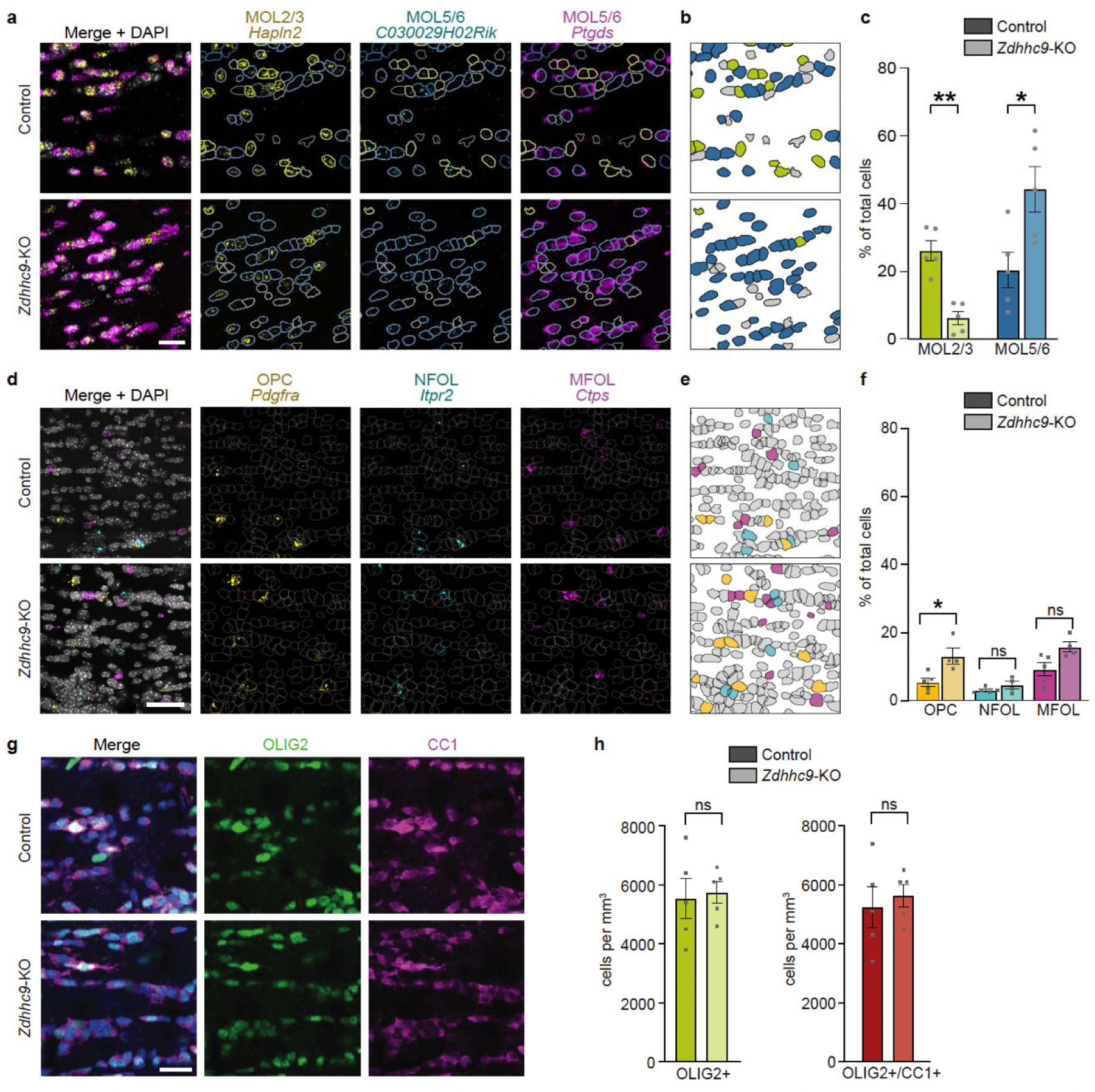
There is a change in mature oligodendrocyte subtype densities, and an increase in the density of OPCs in the corpus callosum of *Zdhhc9*-KO mice. **a** Representative images of fluorescent *in situ* hybridization for MOL2/3 and MOL5/6 (*Hapln2* – MOL2/3; *C030029H02Rik* – MOL5/6; *Ptgds* – MOL5/6) in the P60 corpus callosum (scale bar, 50μm) **b** Representative image of corpus callosum cells colored by oligodendrocyte subtype classification. MOL2/3 cells are colored green; MOL5/6 cells are colored blue; unclassified cells are colored grey. **c** Graph of oligodendrocyte subtype proportions (*n* = 5 mice per genotype). **d** Representative images of fluorescent *in situ* hybridization for immature oligodendrocyte cell-lineage-subtype marker genes (*Pdgfra* -OPCs, *Itpr2* -NFOLs, *Ctps*-MFOLs) in the corpus callosum (scale bar, 50μm). **e** Representative image of corpus callosum cells colored by oligodendrocyte subtype classification. OPC cells are colored yellow; NFOL cells are colored cyan; MFOL cells are colored magenta; unclassified cells are colored grey. **f** Graph of OPC, NFOL and MFOL proportions ( *n* = 4 *Zdhhc9-* KO mice, *n* = 5 control mice). **g** Representative images of OLIG2 (pan oligodendrocyte marker) and CC1 (mature oligodendrocyte marker) immunolabelling in the P60 corpus callosum. **h** Graph of OLIG2+ and OLIG2+/CC1+ proportions (*n* = 5 mice per genotype). Data in **c,f,h** show mean ± SEM. **c**, **f, h** multiple t-tests using Holm-Sidak method, alpha = 0.05.

We next directly quantified whether the density of OPCs or intermediate maturity oligodendrocytes was altered in *Zdhhc9*-KO mice, using a second set of FISH probes including *Pdgfra* (OPC marker), *Itpr2* (NFOL marker) and *Ctps* (MFOL marker). (Figure S4a). *Hapln2* (MOL2/3 marker) was also included in the probe set, to exclude MOL cells from the analysis of immature oligodendrocyte proportions. While we observed no changes in the density of intermediate maturity oligodendrocyte lineage cells, we observed an increase in the density of OPCs, in line with datain Figure 1F, (Figure 2d-f). The increase in OPC density in *Zdhhc9-*KO mice may reflect an intrinsic failure of OPC differentiation or a compensatory increase in OPC proliferation and/or recruitment into the corpus callosum in response to a decreased MOL2/3 population. We also observed no change in the density of oligodendrocyte lineage cells (Olig2^+^) or mature oligodendrocytes (Olig2^+^/CC1^+^) in the corpus callosum of *Zdhhc9-*KO mice (Figure 2g,h). Together, our analysis of differential expression, Bisque deconvolution, and FISH data all corroborate alterations in the subtype proportions of oligodendrocyte lineage cells in the corpus callosum of *Zdhhc9*-KO mice. This includes a reduction in the MOL2/3 subtype, which notably expresses genes linked to myelination, and an increase in OPCs and the MOL5/6 subtype.

### No changes in density of microglia, astrocytes, proliferative or apoptotic oligodendrocytes in *Zdhhc9*-KO mice

Previous studies have ascertained that astrocytes and microglia play an important role in myelination(33–35). We observed no changes in GFAP (marker for astrocyte) or Iba1 (marker for microglia) immunostaining in the corpus callosum, indicating that the observed deficits in myelination in Zdhhc9-KO mice were not caused by increased astrocyte or microglial activity at this time point (Figure S5a,b). We also observed no changes in Olig2/Ki67 and Olig2/CC3 immunostaining demonstrating that differences in oligodendrocyte populations in the corpus callosum is not the result of altered proliferation or apoptosis in oligodendrocytes (Figure S5c-f). Together, these data support the role of *Zdhhc9* in myelination and oligodendrocyte subtype proportions independent of proliferative or apoptotic responses in oligodendrocytes.

### Fewer myelinated axons and myelin compaction abnormalities in the corpus callosum of *Zdhhc9*-KO mice

We next did ultrastructural analyses of the corpus callosum using Transmission Electron Microscopy (TEM) on sagittal brain sections of P60-100 male mice. While we observed no difference in total axon density (myelinated and unmyelinated axons) (Figure 3a,b), there was a significant reduction in the number of myelinated axons in *Zdhhc9* KO mice (Figure 3a,c). This aligns with a reduction in the density of nodes of Ranvier (Figure S6). There was an increase in the diameter of myelinated axons in *Zdhhc9*-KO mice (Figure 3a,d), alongside an increase in the average g-ratio, indicating axonal hypomyelination (Figure 3e,f). Analysis of the ultrastructure of myelinated axons demonstrated a significant increase in the number of axons with myelin abnormalities (Figure 3g, arrowheads in 3a), resembling the residual myelin loops seen in other mouse lines with myelin compaction issues(36). Further testing showed increased distance between intraperiod and major dense lines (increased periodicity), suggesting myelin compaction deficits in *Zdhhc9*-KO mice (Figure 3H). This may partly result from the downregulation of genes such as *Apod* and *Pmp22* in the corpus callosum (Figure 1B,C), which are crucial for myelin compaction(37, 38). Notably, residual myelin loops and uncompacted myelin are typically observed in immature myelin(39, 40). These ultrastructural white matter deficits are similar to those observed in human epilepsy patients(41, 42) and rat models of epileptic seizures(43, 44), suggesting that dysmyelination could affect seizure activity in *Zdhhc9-* KO mice.

**Figure 3.**
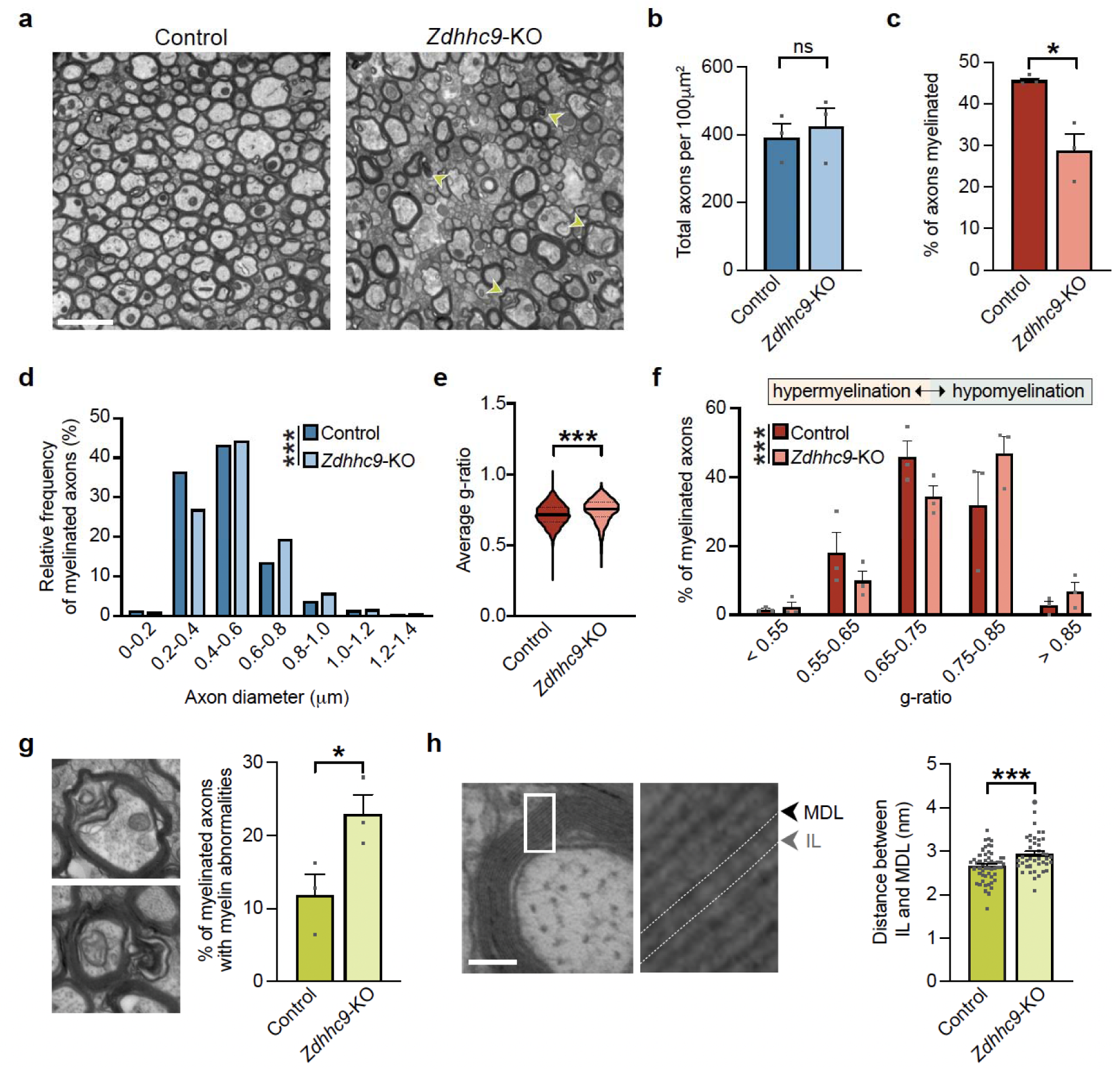
There are fewer myelinated axons and decreased myelin compaction in the corpus callosum of *Zdhhc9*-KO mice. **a** Representative transmission electron microscopy images of the P60 corpus callosum (scale bar, 2mm). **b** Quantification of axon density and **c** the percentage of axons that are myelinated (*n* = 3 mice per genotype). **d** Frequency distribution graph showing the diameter of myelinated axons in the corpus callosum. **e** The average g-ratio of myelinated axons (*n* > 1000 axons from 3 mice per genotype). **f** Frequency distribution graph showing the proportion of myelinated axons with a given g-ratio. **g** Representative images (left) and quantification (right) of axons with myelination decompaction phenotypes (myelin “loops”) (scale bar, 1mm). **h** Representative images of a myelinated axon, showing major dense line (MDL) and intraperiod line (IL) within the myelin sheath (scale bar, 100nm) (left) and quantification of the average distance between IL and MDL (*n* > 30 axons from 3 mice per genotype). Data in **b,c,e,f,g,h** show mean ± SEM. **b,c,g,h** unpaired two-tailed Student’s t-tests; **d,f** Kolmogorov-Smirnov test; **e** Welch’s t-test.

### Decreased expression of integral myelin proteins in the myelin proteome of *Zdhhc9*-KO mice

To better understand how ablation of *Zdhhc9* can impact myelin integrity and compaction, we looked for changes in the myelin proteome using mass spectrometry. We isolated myelin from the whole brain of P23 male mice, an age corresponding with peak myelinogenesis(16) and peak expression of *Zdhhc9*(14) (Figure 4a). Whole brains were used as patients with XLID and *ZDHHC9* LOF mutations exhibit white matter reductions throughout the brain including the corpus callosum, temporal lobes, optic nerve and pituitary stalk(10). 79.6% of all identified proteins were detected in at least 3 of 4 control samples (Dataset S4, Figure S7a), and 77% of all proteins were detected in at least 3 of 4 *Zdhhc9*-KO samples (Dataset S4, Figure S7b). Of these, 90.8% (1864) of all proteins were identified in both genotypes (Dataset S4, Figure S7c). We observed significant differences in the myelin proteome, with an increase in 84 proteins and a decrease in 113 proteins in *Zdhhc9*-KO mice (Figure 4b). Interestingly, GOLGA7, ZDHHC9’s co-factor, was the most significantly decreased protein (Figure 4b). Of the top 15 most abundantly expressed myelin proteins, 4 were significantly decreased in *Zdhhc9*-KO mice (Figure 4c). This included proteins that regulate myelin compaction (CLDN11, MOBP, SIRT2)(45–48), and one protein that has been shown to regulate oligodendrocyte differentiation and the maintenance of mature myelin (MAG)(49). GO term annotation of the 197 differentially expressed proteins showed an association with 25 different biological processes including axon ensheathment, lipid binding, and lipid metabolism (Dataset S5, Figure 4d).

**Figure 4.**
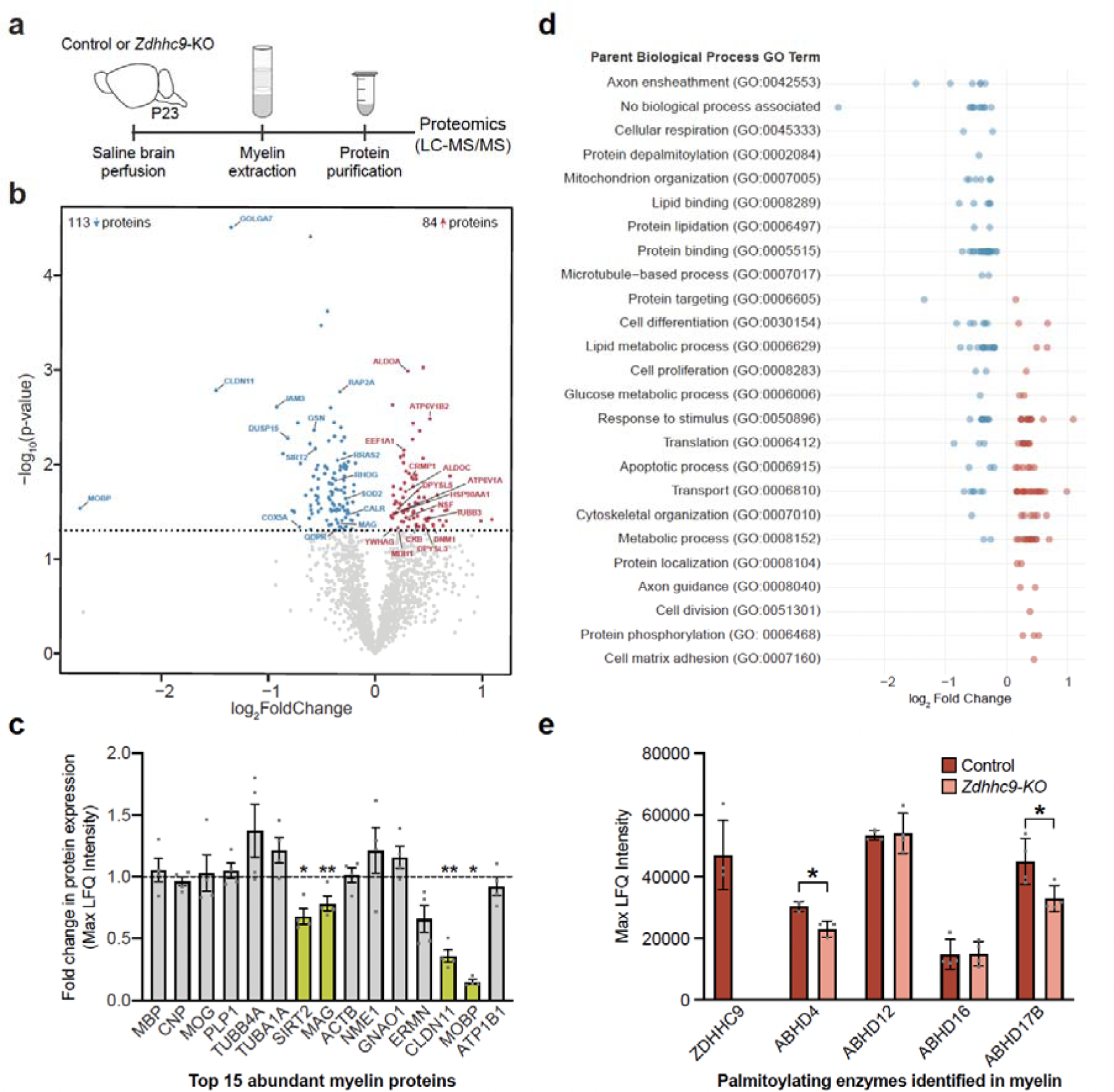
Proteins associated with myelination and myelin compaction are decreased in *Zdhhc9*-KO mice. **a** Illustration showing the pipeline for myelin proteomic analysis of P23 mice. **b** Quantification of fold change in protein expression (*n* = 4 mice per genotype). Each point represents an individual protein; blue denotes proteins that are downregulated, and red denotes proteins that are upregulated in *Zdhhc9*-KO mice relative to controls. The horizontal line denotes the statistical significance threshold required for two tailed t-tests. Labelled proteins denote those that are both differentially expressed and highly abundant (top 10% of control myelin proteome). **c** Graph of Max LFQ Intensity data for the 15 most abundant myelin proteins (*n* = 4 mice per genotype). Normalized expression of each protein relative to control (dotted line). Differentially expressed proteins are highlighted in green. **d** Graph of the biological process GO Term associated with each differentially expressed protein along with their log_2_ Fold Change. **e** Graph of Max LFQ Intensity data for all palmitoylating enzymes identified in myelin (*n* = 4 mice per genotype). All statistical analyses were done using unpaired two-tailed Student’s t-tests.

Interestingly, ZDHHC9 was the only palmitoyl acyl transferase detected in isolated myelin, suggesting a crucial role for this enzyme in regulating the dynamic palmitoylation of myelin proteins within the myelin sheath. Four depalmitoylating enzymes in the ABHD family of palmitoyl thioesterases were identified in control myelin and of these, two (ABHD4 and ABHD17B) were significantly reduced in *Zdhhc9*-KO mice (Figure 4e).

## Discussion

This study highlights an important role for ZDHHC9 in the proper development of oligodendrocyte lineage cells and axon myelination within the corpus callosum. While ablation of *Zdhhc9* does not change the overall density of oligodendrocyte lineage cells, nor the density of mature oligodendrocytes, it results in a greater proportion of MOL5/6 cells and a reduction in MOL2/3 cells. As MOL2/3 cells are particularly enriched for genes associated with myelination, the loss of MOL2/3 cells could in part explain the reduction in the number of myelinated axons. Loss of *Zdhhc9* also results in ultrastructural changes in myelin integrity likely driven by a decrease in key myelin proteins such as MAG, CLDN11, and MOBP. This work suggests that disruptions in oligodendrocyte differentiation and axon myelination could be integral to the pathophysiology observed in patients with XLID associated with *ZDHHC9* loss-of-function mutations. Previous studies have identified several ZDHHC enzymes that can modulate gene expression through the palmitoylation of transcription factors(50–53), chromatin remodelers(54–56), and histones(57, 58). Thus, altered gene expression observed in the corpus callosum of *Zdhhc9*-KO mice could be a result of changes in the palmitoylation of these substrates as well as signaling pathways that alter gene transcription. Notably, in our dataset we identified altered expression of several genes encoding proteins that further regulate gene expression. For example, we observed decreased expression of *Insig1*, a modulator of sterol regulatory element-binding proteins (SREBPs)(59), which are key regulatory transcription factors for lipid metabolism and synthesis(60). The decreased expression we observed in a large cluster of genes involved in cholesterol metabolism is particularly interesting given that cholesterol is the only essential lipid in myelin membranes(61) and is known to regulate the extent of myelination(62, 63), influence myelin compaction(64), and acts as a signaling molecule to drive oligodendrocyte differentiation and maturation(65).

Numerous studies in recent years have set out to identify the function of the numerous transcriptionally distinct oligodendrocyte subtypes that have been discovered using scRNAseq(15, 32). Here, we leveraged selective markers spanning all stages of oligodendrocyte maturation and observed a selective decrease in *Hapln2*+ MOL2/3s and an increase in both *Pdgfra*+ OPCs and *Ptgds+/C030029H02Rik+* MOL5/6s. *Zdhhc9* is most highly expressed in intermediate maturity oligodendrocytes (MFOLs), and expression remains elevated in mature oligodendrocytes(ADD MARQUES REF). It is therefore possible that oligodendrocytes maturation to *Hapln2*+ MOLs is impaired in the absence of *Zdhhc9*. Although altered proportions in the MOL subtypes have been observed in multiple disease states(66, 67),the precise function of each of the transcriptomically classified subtypes of MOL remains unclear. It is however noteworthy that the MOL2/3 population that are depleted in *Zdhhc9* KO mice are distinguished by their elevated expression of myelin-related genes, compared with other MOL subtypes(67). Future work is needed to determine precisely how changes in the proportions of MOL subtypes in *Zdhhc9* KO mice contribute to the myelin deficits observed.

Our FISH and RNAseq analyses also revealed increased OPC populations in the corpus callosum of *Zdhhc9-*KO mice. While expression of *Zdhhc9* in OPCs is low, our proteomic analysis revealed that a key regulator of OPC differentiation, SIRT2, was downregulated in *Zdhhc9-*KO mice(68). The failure of OPCs to differentiate could therefore be responsible for the increased OPC density in *Zdhhc9-*KO mice. Alternatively, the increase in OPC density could reflect a compensatory mechanism to repopulate depleted MOL2/3 oligodendrocytes and/or a response to the decrease in myelinated axons(69). These observations nevertheless suggest that the balance between mature oligodendrocytes and their progenitors is disrupted in *Zdhhc9-*KO mice. This disruption could be a common feature of intellectual disabilities, as impairments in other intellectual disability risk genes, such as PAK3(70) and ATRX(71), have been shown to increase OPC densities, leading to compromised myelination.

We also observed ultrastructural changes in myelin integrity in the corpus callosum and changes in the composition of the myelin proteome in *Zdhhc9*-KO mice that may underlie functional deficits in patients with *ZDHHC9*-XLID. Interestingly, MAG and SIRT2, two highly abundant myelin proteins that regulate myelin compaction(72, 73) are decreased in *Zdhhc9-*KO myelin. Additionally, proteins such as MOBP and CLDN11, which are enriched in compact myelin(74), were less abundant in *Zdhhc9*-KO myelin, supporting our ultrastructural data demonstrating decreased myelin compaction. Alterations in the myelin proteome could be the result of altered gene expression, protein stability and/or disrupted targeting and sorting of protein into the myelin membrane. Indeed, palmitoylation has been shown to be required for trafficking key myelin proteins into the myelin sheath(75) and previous work from our lab has identified key myelin proteins including MOBP and PLP to be substrates for ZDHHC9(14). Overall, we propose a model for decreased myelination in the corpus callosum of *Zdhhc9* KO mice that is a result of i) changes in the proportions of oligodendrocyte lineage cells and ii) decreased expression of numerous proteins that are involved in cholesterol biosynthesis and maintaining myelin integrity. The findings of this study establish a critical role for ZDHHC9 in oligodendrocyte development and myelination and identify dysregulated signaling pathways that may underlie the decrease in white matter volume, epileptic comorbidities and intellectual disability seen in patients with *ZDHHC9* mutations.

## Materials and Methods

### Animals

*Zdhhc9*-KO mice (B6;129S5-Zdhhc9tm1Lex/Mmucd) were originally obtained from the Mutant Mouse Resource and Research Center (MMRRC, University of California Davis). A mutation in *Zdhhc9* was introduced through homologous recombination, targeting coding exon 1 (NCBI ascension: NM_172465.1). All experiments were performed on mice between 2-4 months of age, except for analysis of the myelin proteome, where mice were 23 days of age. Mice were housed in same-sex groups of 2-5 within optimouse cages, with bedding, a plastic hut and nesting material for enrichment. Food and water were available *ad-libitum*. Ear notches were collected at weaning (21 days of age) for identification, and genotyped by Transnetyx (Cordova, TN). All procedures followed guidelines from the Canadian Council on Animal Care and were approved by the UBC Animal Care Committee.

### Fluoromyelin staining and analysis

Thickness and area measurements were obtained using across six regions (each 350-600um in length) spanning the anterior-posterior axis of the corpus callosum. Measurements were obtained from within each of the Genu (1.1 to 0.26), Body (-0.01 to -1.34) and Splenium (**-**1.45 to -2.54) of the corpus callosum, based on stereotaxic coordinates in Franklin and Paxinos(76). For each mouse, 2-5 sections were sampled from each region, and measures were obtained either at midline, or averaged bilaterally across cerebral hemispheres.

### Tissue collection for RNA-sequencing

Mice were anesthetized under isoflurane (5%, 1.5L/min), and decapitated. Brains and optic nerves were dissected, and flash frozen in pre-chilled isopentane at -52**°**C for 30 seconds. Brains were then stored at -80°C until sectioning. The corpus callosum (Bregma 0.98mm – Bregma 0.62mm) was sectioned coronally at 100μm on a cryostat and mounted onto glass slides. Slides were then stored at -80°C until used. Corpus callosum samples were dissected from frozen brain sections, while being cooled on dry ice. RNA was extracted from samples using the Total RNA Purification Micro Kit (Cat. 35300, Norgen Biotek Corp) following the supplied protocol. Samples were then kept at -80**°**C until analyzed by the RNA sequencing core at the School of Biomedical Engineering Sequencing Core at the Biomedical Research Center (University of British Columbia).

### RNA-sequencing

Sample quality control was performed using the Agilent 2100 Bioanalyzer or the Agilent 4200 Tapestation. Qualifying samples were then prepped following the standard protocol for the Illumina Stranded mRNA prep (Illumina). Sequencing was performed on the Illumina NextSeq2000 with Paired End 59bp × 59bp reads.

### RNA-sequencing analysis

RNA-seq raw sequencing reads were processed with kallisto(77) using the reference transcriptome for mm10 downloaded from Ensembl (www.ensembl.org) and the resulting counts for different isoforms were summed up to obtain counts at the gene level for each sample. The differential expression analysis was then performed with DESeq2(78) and genes achieving the significance cut-off of FDR < 0.05 were considered significantly differentially expressed. Marker genes for each cell type were extracted from the file l5_all.agg.loom downloaded from the Mousebrain website (http://mousebrain.org/adolescent/downloads.html).

### Quantitative PCR

Total RNA was extracted from samples using the Total RNA Purification Micro Kit (Cat. 35300, Norgen Biotek Corp) following the supplied protocol. Quantitative PCR with reverse transcription was performed on a ViiA 7 instrument (Thermo Fisher Scientific) using SYBR Green master mix (Thermo Fisher Scientific). The relative amounts of the messenger RNAs studied were determined by means of the 2−ΔΔ.CT method, with Gapdh as the reference gene and the control genotype as the invariant control.

### Bioinformatics

Biological process enrichment analysis was done using PANTHER overrepresentation test (Released 2022-10-13). Functional interaction networks of the differentially expressed genes were investigated using the Search Tool for the Retrieval of Interacting Genes (STRING) 11.5(79). A threshold confidence level of 0.4 was used to identify protein interactions and seven types of protein interactions were used for network generation, including neighborhood, gene fusion, co-occurrence, co-expression, experimental, database knowledge, and text mining. The Mouse Genome Informatics (RRID:SCR_006460) database(80) was used to assign GO classifications for the list of differentially expressed genes and differentially expressed proteins. Cell type proportion estimates were performed using Bisque deconvolution(81) of our RNAseq data using the Brain Initiative Cell Census Network 2.0(32) as the reference scRNAseq dataset. Clusters from Yao et al., 2023 were mapped onto the widely adopted nomenclature(15, 18, 66, 82, 83) used for oligodendrocyte lineage first defined by Marques et al., 2016 using the expression of marker genes in each cluster: clusters 5266-5271 = OPC, 5272-5277 = COP, 5280 = NFOL1, 5281 = NFOL2, 5283 = MFOL1, 5282 = MFOL2, 5288 = MOL1, 5284 & 5286 = MOL2/3, 5285 & 5287 = MOL5/6.

### Fluorescent in situ hybridization

Fluorescent in situ hybridization was performed with target retrieval, pretreatment, hybridization, amplification, and detection according to the RNAscope Multiplex Fluorescent Reagent Kit v2 Assay User Manual for Fixed Frozen Tissue (ACD). Multiplexed FISH probes for *Pdgfra* (480661-C2), *Itpr2* (579391-C3), *Ctps* (1149611-C4), *Wfdc18* (430051-C2), *Hapln2* (1149621-C1), *Ptgds* (492781-C4), and *C030029H02Rik* (1301981-C3) were used as marker gene probes to identify OPCs, NFOLs, MFOLs, MFOL2, MOL2/3, and MOL5/6 respectively. *Wfdc18* was used as a marker to verify cells were not MFOLs. *Hapln2* was used with *Pdgfra, Itpr2,* and *Ctps* to verify cell assignment as an immature oligodendrocyte subtype. Slides were cover slipped with Prolong Gold Antifade Mountant (Invitrogen, Cat. P36930). Once dried, slides were kept at 4**°**C until imaged.

### Fluorescent in situ hybridization analysis

First, a region of interest was created around the corpus callosum using the polygon tool in FIJI. Cell ROIs within the corpus callosum were then generated from the DAPI channel using a custom FIJI macro. In short, the macro consisted of auto thresholding the image, converting the image to a mask, applying Gaussian blur (sigma = 3), binarizing the image, inverting the image, applying Watershed, and dilating particles by 1um. Finally, only nuclei between 50-310μm^2^ and a circularity of 0.5-1.0 were selected for analysis. *Pdgrfa*, *Itpr2*, and *Ctps* channels were then each individually thresholded and percent coverage for each probe was measured in each cell ROI. Cells were assigned as OPC, NFOL, or MFOL according to the marker probe with the highest percent coverage in each cell nucleus. Cells were only counted that had a dominant probe percentage coverage > 4% (minimum marker gene expression).

For MOL2/3 and MOL5/6 analysis, a similar region of interest was created around the corpus callosum in FIJI. Nuclei were then identified and segmented from the maximum intensity projection image using the IdentifyPrimaryObjects module in CellProfiler v4.2.6(84). Nuclei were then dilated by 3 pixels to capture near-nuclear RNAscope staining. A label image of the segmented nuclei was identified and used to make nuclear ROIs in FIJI through the “Label Image to ROIs” plugin from the PTBIOP update site. RNAscope channels were manually thesholded as above and percent coverage per cell measured. Cell assignment was carried out in R v4.3.2 using RStudio v2023.12.1.402 (Posit Software) as described previously(82). Briefly, cells were assigned to MOL2/3 if their *Hapln2* coverage exceeded an empirical threshold (16%) of the nuclear area and if their *Hapln2* coverage was greater than the *Hapln2* area was greater than the *Ptgds* position. Cells were assigned to MOL5/6 if their *Ptgds* coverage was greater than 7% of nuclear area. As *Ptgds* alone does not mark MOL5/6 exclusively (Figure S4a), we additionally filtered this class of cells for inclusion of the long non-coding RNA *C030029H02Rik* and exclusion of the MFOL2 marker *Wfdc18*.

### Immunostaining

All immunostaining was completed on free-floating sections. Sections were first washed in PBS (0.1M, pH 7.4), 3 times for 5 min. Antigen retrieval was then completed by incubating sections in a sodium citrate buffer (10mM, pH 6.0) with Tween 20 (0.5%) for 20min at 80°C. Sections were then cooled and were incubated in PBS with 0.1% triton-X (PBS-TX) and 10% goat serum for 3 hours to reduce subsequent non-specific anti-body binding. The sections were then incubated in PBS-TX with 5% goat serum containing primary antibodies for Na_v_1.6 (1:500 dilution, ASC-009, Alamone); or Olig2 (1:500, NBP1-28667, Novus) and CC-1 (1:100, ab16794, Abcam); or GFAP (1:500, ab7260, Abcam) and Iba1 (1:500, ab178846, Abcam); or Olig2 (1:500, NBP1-28667, Novus) and Ki67 (1:500, ab15580, Abcam); or Olig2 (1:500, NBP1-28667, Novus) and CC3 (1:500, 9661S, Cell Signaling Techno) were incubated overnight at 4**°**C. Following primary antibody incubation, sections were washed in PBS-TX (3 x 5min) and incubated in PBS-TX containing secondary goat anti-rabbit AlexaFluor 488 (1:500 dilution, Ab150077, Abcam) or goat anti-mouse AlexaFluor 488 (1:500, A-21141, Invitrogen) and goat anti-rabbit AlexaFluor 647 (1:500, A-21244, Invitrogen). Sections were then washed in PBS-TX (3 x 5min) and counter-stained with DAPI (1:500) in PBS-TX for 10 minutes. Lastly, sections were washed in PBS (3 x 5min), mounted onto Superfrost Plus glass slides (Fisher Scientific), and coverslips applied using the ProLong Gold anti-fade (Invitrogen) mounting medium.

### Electron microscopy sample preparation

Brains were first sectioned coronally on a vibratome (Leica, VT1000) at 200μm thickness. Comparable posterior sections (-1.82 to -2.30 from Bregma) were selected for each animal, and sections were washed three times in sodium cacodylate buffer (0.1M). The hemispheres of each section were bisected using a razor blade, and a brain punch (0.50mm, Ted Pella) was used to make a fiducial hole dorsal to the corpus callosum. Samples were then post-fixed with 1% osmium tetroxide and 1.5% potassium ferricyanide, stained *en bloc* with 2% uranyl acetate, dehydrated in an alcohol series, and finally embedded in Epon 812 resin (Electron Microscopy Sciences). 70nm ultra-thin sagittal sections were obtained along the medial edge of the corpus callosum using an ultramicrotome (UC7, Leica) and mounted onto 2×1 mm Formvar-coated copper slots (Electron Microscopy Sciences). Ultra-thin sections were then grid stained with 2% uranyl acetate for 12 minutes and Reynolds’ lead citrate stain for 6 minutes. Slots were imaged on a FEI Tecnai Spirit transmission electron microscope at 80 kV. Images of the corpus callosum were taken in the dorsal region of the corpus callosum.

### Electron microscopy analysis

For each animal 1-4 images taken at 9300X magnification (105.3μm^2^), ∼100-300 axons per image, were analyzed using software for semi-automatic myelin *g*-ratio quantification (Myeltracer(85)). Myelinated axons with residual myelin loops and splitting of myelin were classified as axons with myelin abnormalities. To assess myelin decompaction, 10-15 higher magnification images were taken at 68000X magnification and ∼20 axons per animal were analyzed.

### Microscopy

All myelin markers, nodal markers, and fluorescent in situ hybridization images were obtained using a Zeiss LSM 880 confocal microscope equipped with 405, 488, 543, 594, and 633 laser lines, 3 multialkali spectral PMTs and an Airyscan detector. Images of fluoromyelin were obtained at a single optical plane, using a 20x/0.8 NA objective with tile scans. Images of fluorescent in situ hybridization were acquired with at 63x/1.4 NA oil objective and were comprised of *Z*-stacks taken at 1.00 μm intervals through the entire section. Immunohistochemistry images were acquired using the Airyscan detector in the FastAiry mode. Tile scans covering the medial corpus callosum were acquired, maximum intensity *Z-*projected, and stitched in Zen Black software (Zeiss). All images were obtained using optimized laser settings for all five channels that were identical between sections.

### Myelin purification

A myelin-enriched membrane fraction was biochemically purified from mouse brains by sucrose density centrifugation and osmotic shocks as described previously(86, 87). Briefly, flash frozen mouse brains were thawed in 0.32M sucrose supplemented with protease inhibitor cocktail tablets (cOmplete Mini EDTA-free; Roche, Mannheim, Germany). Thawed brains were next manually homogenized using a pestle and 16G/25G needle. Brain homogenate was then layered carefully on top of 0.85M sucrose and centrifuged. Following the first sucrose gradient, the band of myelin-enriched fraction found at the 0.32M/0.85M sucrose interphase was collected and washed with distilled water. Two osmotic shocks using cold water were used to diminish cytoplasm and microsomes, followed by a second sucrose gradient and washing. To delipidate the purified myelin protein samples underwent a series of methanol, chloroform, and distilled water washes.

### Liquid-Chromatography-Tandem Mass-Spectrometry (LC-MS/MS)

Myelin protein samples, boiled in SDS sample buffer, were run on 10% SDS/PAGE gel. Proteins were detected with colloidal Coomassie G-250 and digested out of the gel as described previously(88). Samples were purified by solid phase extraction on C-18 STop And Go Extraction (STAGE) Tips. Samples were separated using a NanoElute UHPLC system (Bruker Daltonics) with Aurora Series Gen2 (CSI) analytical column (25cm x 75μm 1.6μm FSC C18, with Gen2 nanoZero and CSI fitting; Ion Opticks, Parkville, Victoria, Australia) heated to 50°C (by Column toaster M, Bruker Daltonics) and coupled to timsTOF Pro (Bruker Daltonics) operated in DDA-PASEF mode. In a standard 60 min run, the gradient was from 2% B to 12% B over 30 min, then to 33% B from 30 to 60 min, then to 95% B over 0.5 min, held at 95% B for 7.72 min. Before each run, the analytical column was conditioned with 4 column volumes of buffer A. Where buffer A consisted of 0.1% aqueous formic acid and 0.5 % acetonitrile in water, and buffer B consisted of 0.1% formic acid in 99.4 % acetonitrile. The NanoElute thermostat temperature was set at 7°C. The analysis was performed at 0.3 μL/min flow rate.

The Trapped Ion Mobility – Time of Flight Mass Spectrometer (TimsTOF Pro; Bruker Daltonics, Germany) was set to Parallel Accumulation-Serial Fragmentation (PASEF) scan mode for DDA acquisition scanning 100 – 1700 m/z with 5 PASEF ramps. The capillary voltage was set to 1800V, drying gas to 3L/min, and drying temperature to 180°C. The MS and MS/MS spectra were acquired from m/z 100 – 1700. As for TIMS setting, ion mobility range (1/k0) was set to 0.70 – 1.35 V·s/cm^2^, 100ms ramp time and accumulation time (100% duty cycle), and ramp rate of 9.42Hz; this resulted in 0.64s of total cycle time.

Linear precursor repetitions were applied with a target intensity of 21,000 and 2500 intensity threshold. The active exclusion was enabled with a 0.4 min release. The collision energy was ramped linearly as a function of mobility from 27eV at 1/k0 = 0.7 V·s/cm^2^ to 55eV at 1/k0 = 1.35 V·s/cm^2^. Isolation widths were set at 2.07 m/z at < 400 m/z and 3.46 m/z at > 1000 m/z. Mass accuracy: error of mass measurement is typically within 3 ppm and is not allowed to exceed 7 ppm. For calibration of ion mobility dimension, the ions of Agilent ESI-Low Tuning Mix ions were selected (m/z [Th], 1/k0 [Th]: 622.0290, 0.9915; 922.0098, 1.1986; 1221.9906, 1.3934).

TimsTOF Pro was run with timsControl v. 3.0.0 (Bruker). LC and MS were controlled with HyStar 6.0 (6.0.30.0, Bruker).

### Protein Search and Analysis

Data was analyzed with Fragpipe V18.0, MSFragger V3.5, Philosopher V4.4.0 (89) and quantitation performed using the MaxLFQ algorithm with match between runs enabled(90, 91). The searches were matched to a mus musculus database downloaded from uniprot.org (reviewed sequences; RRID:SCR_002380) and a contaminant and decoy database. Default search parameters were used except for the following: 50 ppm mass tolerances were used for each of the MS and MS/MS, and MSMS not required for LFQ. Accurately identified peptides were only confined to those with IonScores higher than 99% confidence. Contaminant and reverse hits were removed.

In LFQ, the intensities of the thus recorded peaks are taken as proxies for peptide abundance. With fragmentation spectra (MS2 spectra) used for peptide identification. In LFQ, each sample was separately analyzed on the mass spectrometer, and differential expression was obtained by comparing relative intensities between runs for the same identified peptide(92). Log ratios were calculated as the difference in average log_2_ LFQ intensity values between control and *Zdhhc9*-KO groups.

### Statistical analyses

Prior to analysis, normality and homogeneity of variance were tested for. All values in graphs report the mean ± standard error of the mean. Fluoromyelin staining, FISH, TEM, nodal staining, and proteomics data was analyzed with unpaired two-tailed Student’s t test t-tests (p <0.05). TEM axon size and g-ratio distribution analyses were completed using the Kolmogorov–Smirnov test. All major statistical analyses, outlier analyses and graph generation were conducted using GraphPad Prism (Version 8.4.3) software (GraphPad, CA, United States). Asterisks and hashtags are used to denote levels of statistical significance within all graphs (*p < 0.05; **p<0.01; ***p<0.001).

## Acknowledgments

The authors thank Drs. Gareth Thomas and Shin Hyeok Kang (Temple University) for insightful discussions throughout the study. We thank the UBC Proteomic Core as well as Dr. Naoji Yubuki, the UBC Bioimaging Facility (RRID: SCR_021304) and the UBC School of Biomedical Engineering Sequencing Core at the Biomedical Research Center for technical help. Dr. Mark Cembrowski’s lab (UBC) provided expertise on FISH. This work is supported by grants from CIHR (FDN-159907) (S.X.B.) and NIH (AWD-020422) (G.T., S.H.K., & S.X.B.), as well as by computational resources made available through the NeuroImaging and NeuroComputation Centre at the Djavad Mowafaghian Centre for Brain Health (RRID:SCR_019086) and the Dynamic Brain Circuits in Health and Disease Research Excellence Cluster DataBinge Forum.

## Figures

**Fig. S1.**
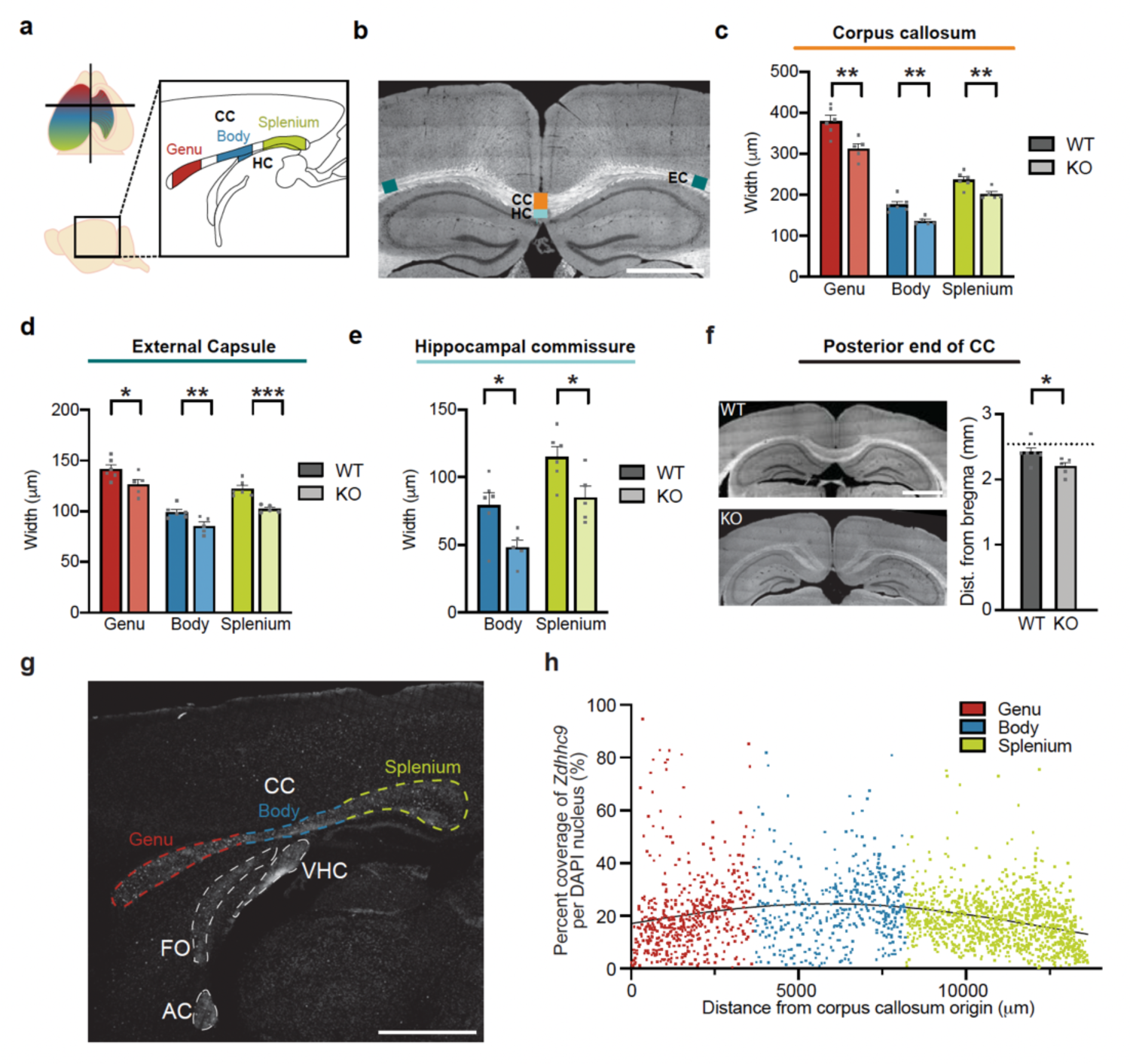
There is a uniform reduction in the width of white matter tracts in Zdhhc9-KO mice and a uniform expression of Zdhhc9 throughout the anterior posterior axis of the corpus callosum. **a** (Top left) Schematic of the brain (horizontal plane), illustrating the spatial relationship between the position of cortical regions and the respective location their projections cross the anterior posterior axis of the corpus callosum. (Right) Schematic of the anterior-posterior axis of the corpus callosum, colored based on its main anatomical subdivisions: genu (red), body (blue), splenium (green). The location of these colored regions indicates where measurements were taken for subsequent morphological analyses. **b** Representative image illustrating white matter tracts that were quantified (corpus callosum, hippocampal commissure, external capsule), and the relative position where measurements were taken within each tract (scale bar, 1mm). **c** Width of the corpus callosum **d** the external capsule and **e** the hippocampal commissure in control and *Zdhhc9*-KO mice. (c,d,e *n =* 5 *Zdhhc9-*KO mice, *n* = 6 control mice) **f** Representative images of the posterior location at which the corpus callosum terminates in control and *Zdhhc9*-KO mice (scale bar, 1mm). Distance from bregma at which the corpus callosum terminates in control and *Zdhhc9*-KO mice. **g** Representative image of FISH for *Zdhhc9* along the anterior-posterior axis of the corpus callosum (scale bar, 1mm). White matter tracts are labeled with dashed lines, along with the main subdivisions of the corpus callosum (colored as in **a**). **h** Percent coverage of *Zdhhc9* labeling per cell as a function of distance from the origin of the corpus callosum. Points are colored based on sub-regions delineated in **h** (*n* of at least 500 cells per region from 1 animal). Data in **c,d,e,f** represent mean + SEM, all statistical tests were two-tailed unpaired Student’s t-tests. CC=corpus callosum, HC=hippocampal commissure, FO=Fornix, VHC=Ventral hippocampal commissure, AC=anterior commissure.

**Fig. S2.**
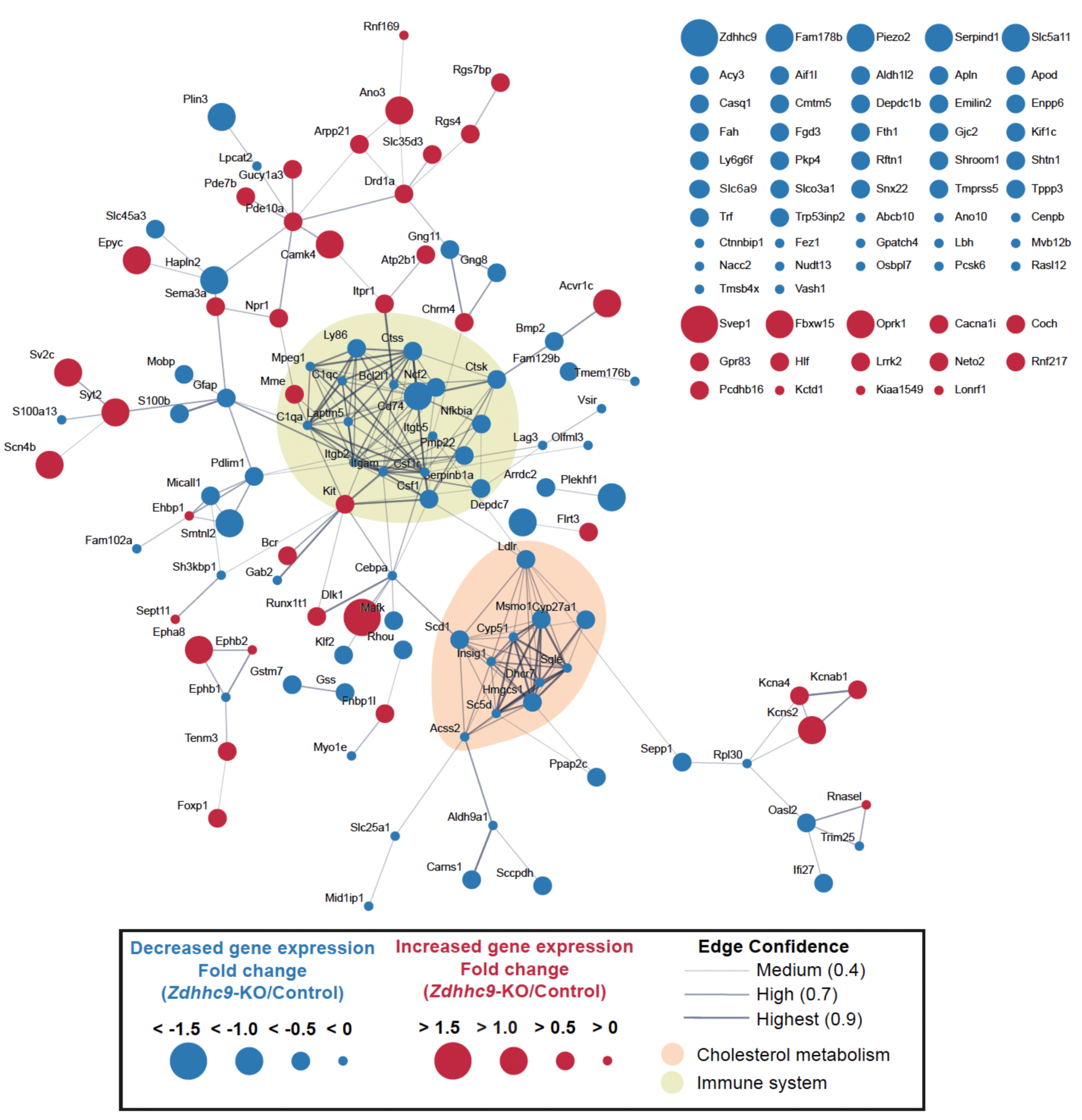
Protein-protein interaction analysis of differentially expressed genes. STRING analysis highlighting protein-protein interactions between differentially expressed genes identified from RNAseq (left). Blue nodes represent downregulated genes, red nodes represent upregulated genes and the node size depicts the magnitude of the fold change in gene expression. Lines (edges) between nodes show predicted protein-protein interactions, and the corresponding line-width reflects the confidence of protein-protein interactions. Two groups of proteins with high densities of interactions were noted and contain proteins with known roles in cholesterol metabolism (orange circle) and immune system function (green circle). Differentially expressed genes without predicted protein-protein interactions are also shown (right).

**Fig. S3.**
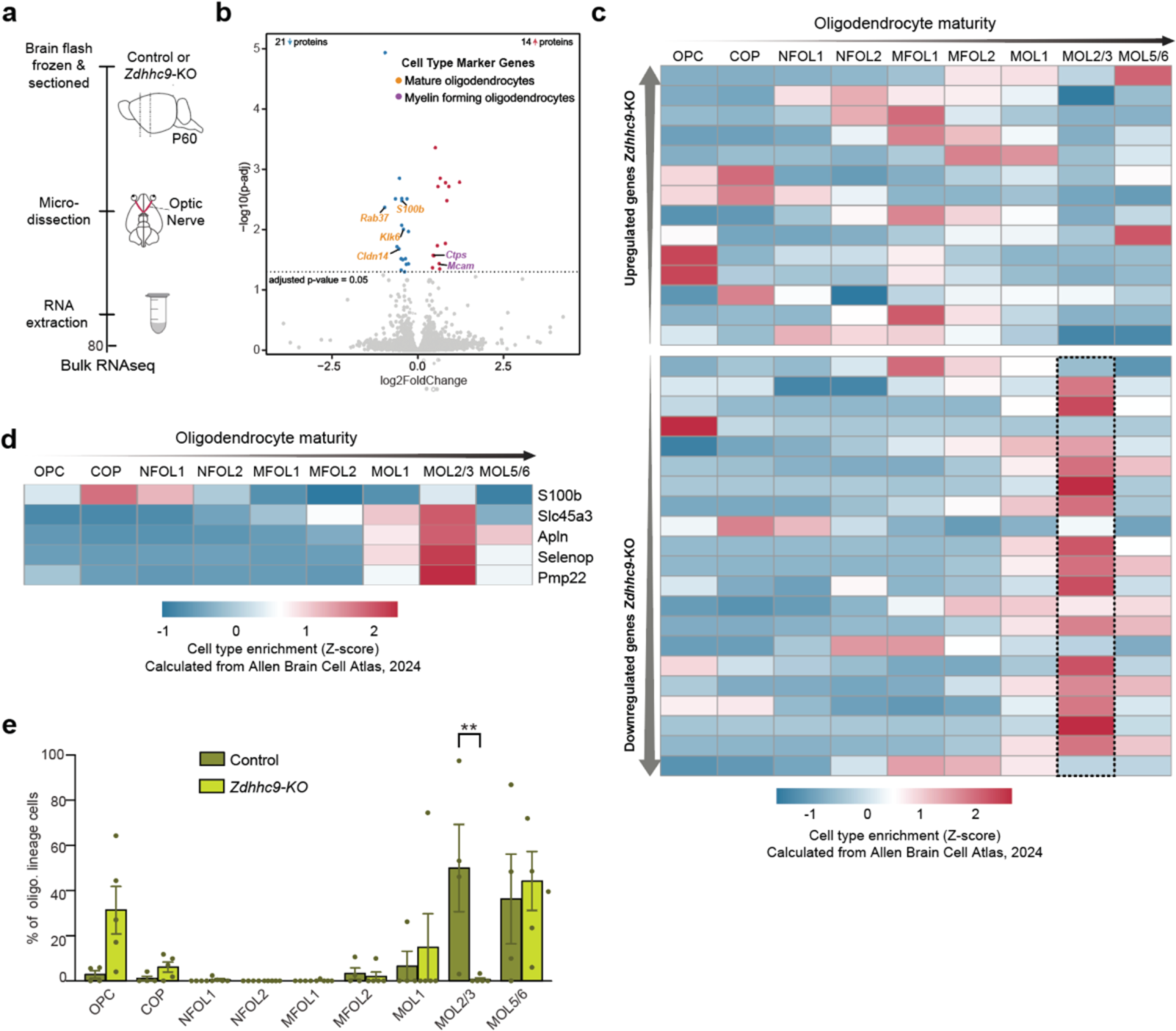
There is change in gene expression in the optic nerve of *Zdhhc9*-KO mice. **a** Illustration showing pipeline for RNAseq analysis in P60 mice. **b** Quantification of fold change in gene expression in *Zdhhc9*-KO optic nerve compared to control mice (*n* = 5 *Zdhhc9*-KO, *n* = 4 control mice). Each point represents an individual gene; blue denotes genes that are downregulated, and red denotes genes that are upregulated in *Zdhhc9*-KO mice relative to controls. The horizontal line denotes the statistical significance threshold required for two tailed t-tests; p-values were adjusted to account for multiple comparisons. Top 5 downregulated and upregulated genes are annotated, along with genes differentially expressed in both corpus callosum and optic nerve. **c** Heatmap showing the expression of the 35 differentially expressed genes within different oligodendrocyte subtypes using scRNAseq data from ABC Atlas. Genes are listed (top to bottom) by decreasing log2 Fold Change (i.e. from most upregulated to the most downregulated). Dotted black boxes highlight overall enrichment of upregulated and downregulated genes. **d** Heatmap showing the expression of genes differentially expressed in both corpus callosum and optic nerve within the different oligodendrocyte subtypes. **e** Bisque bioinformatic estimation of oligodendrocyte cell type proportions in the optic nerve, performed on bulk RNAseq data from this study and using Allen brain ABC atlas as a reference scRNAseq dataset.

**Fig. S4.**
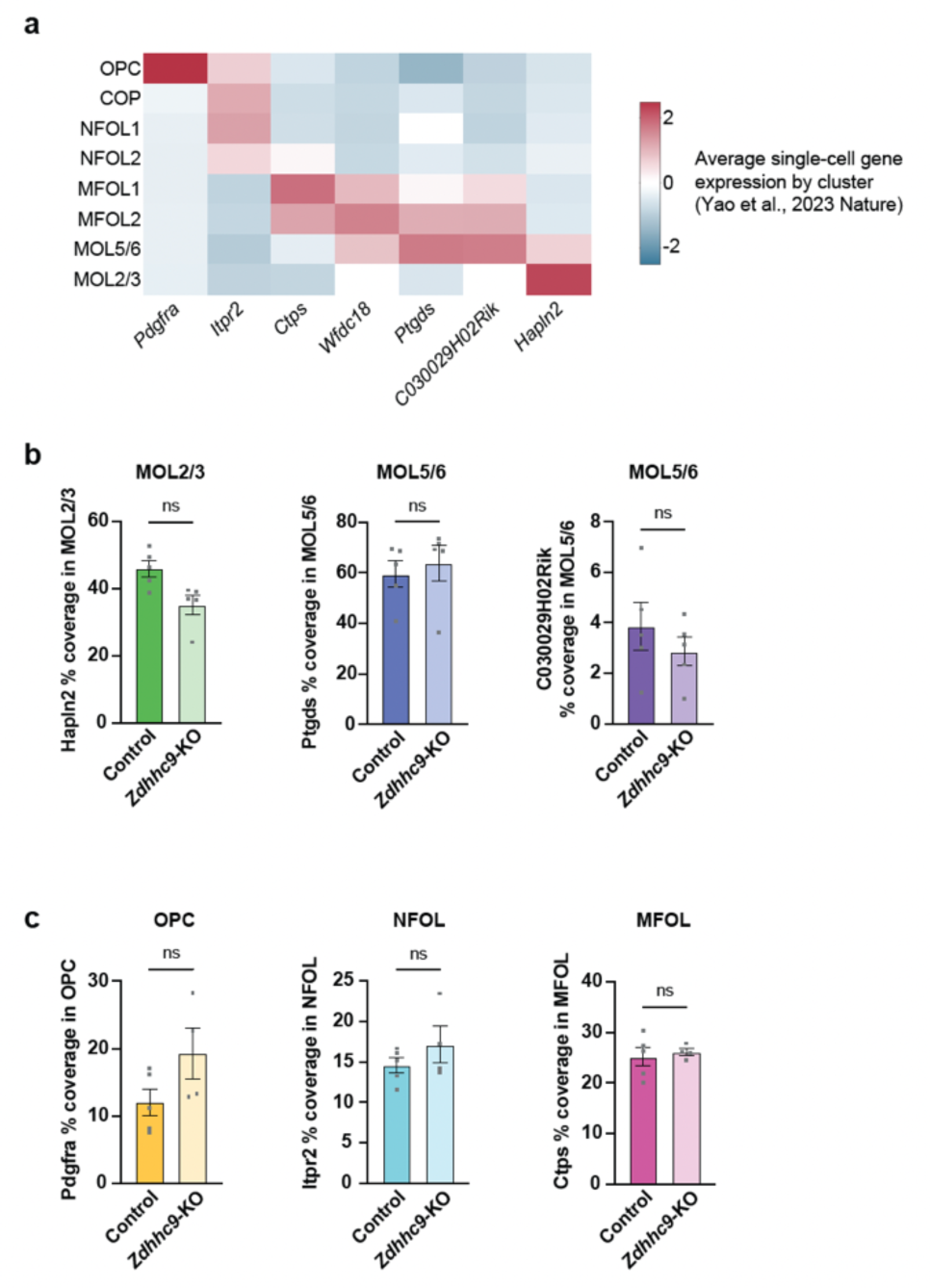
Overall levels of oligodendrocyte lineage marker gene expression does not differ between *Zdhhc9*-KO and control mice. **a** Heatmap of gene expression for our oligodendrocyte lineage marker genes, across different cell types and maturation stages within the oligodendrocyte lineage (scRNAseq from ABC Atlas). **b,c** Graphs depicting the average coverage of oligodendrocyte lineage probes per cell, (*n =* 5 mice per genotype, **b**; *n* = 4 *Zdhhc9*-KO mice, *n* = 5 control mice, **c**). Data shows mean + SEM, and statistical analyses were done using unpaired two-tailed Student’s t-tests.

**Fig. S5.**
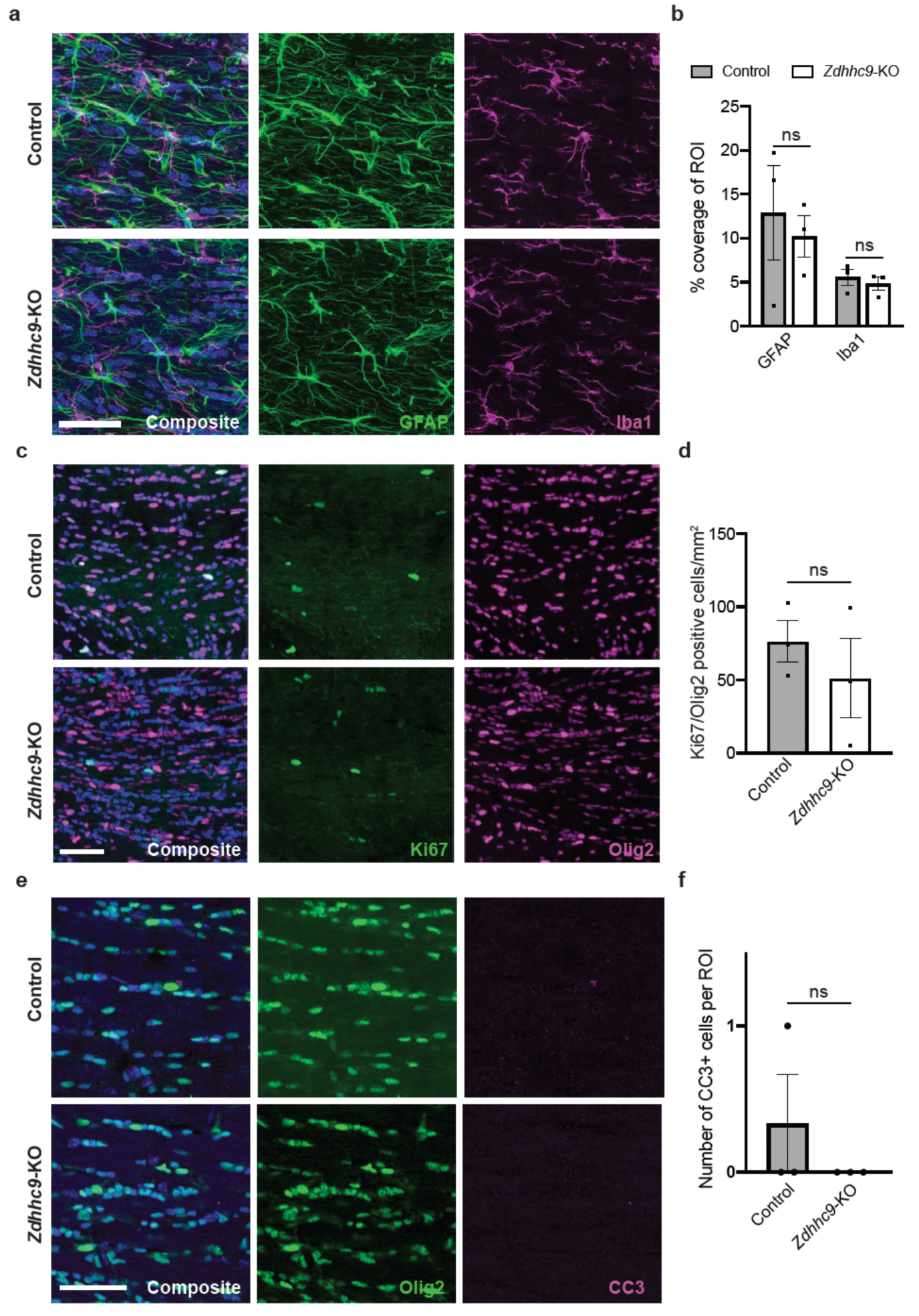
There are no differences in the density of microglia, astrocytes, proliferative or apoptotic oligodendrocytes in *Zdhhc9*-KO mice. **a** Representative images of GFAP (astrocytes) and Iba1 (microglia) immunostaining in the P60 corpus callosum. **b** Quantification of the percent coverage of each marker within the ROI. (*n* = 3 mice per genotype). **c** Representative images of Ki67 (proliferating cells) and Olig2 (oligodendrocytes) immunostaining in the corpus callosum. **d** Quantification of proliferating oligodendrocytes (Ki67+/Olig2+ cells) in the corpus callosum. (n = 3 mice per genotype) **e** Representative images of Olig2 (oligodendrocytes) and CC3 (apoptotic cells) immunostaining in the corpus callosum. **f** Quantification of apoptotic oligodendrocytes. (*n* = 3 mice per genotype). Data in **b,d,f** show mean ± SEM, unpaired two-tailed Student’s t-tests.

**Fig. S6.**
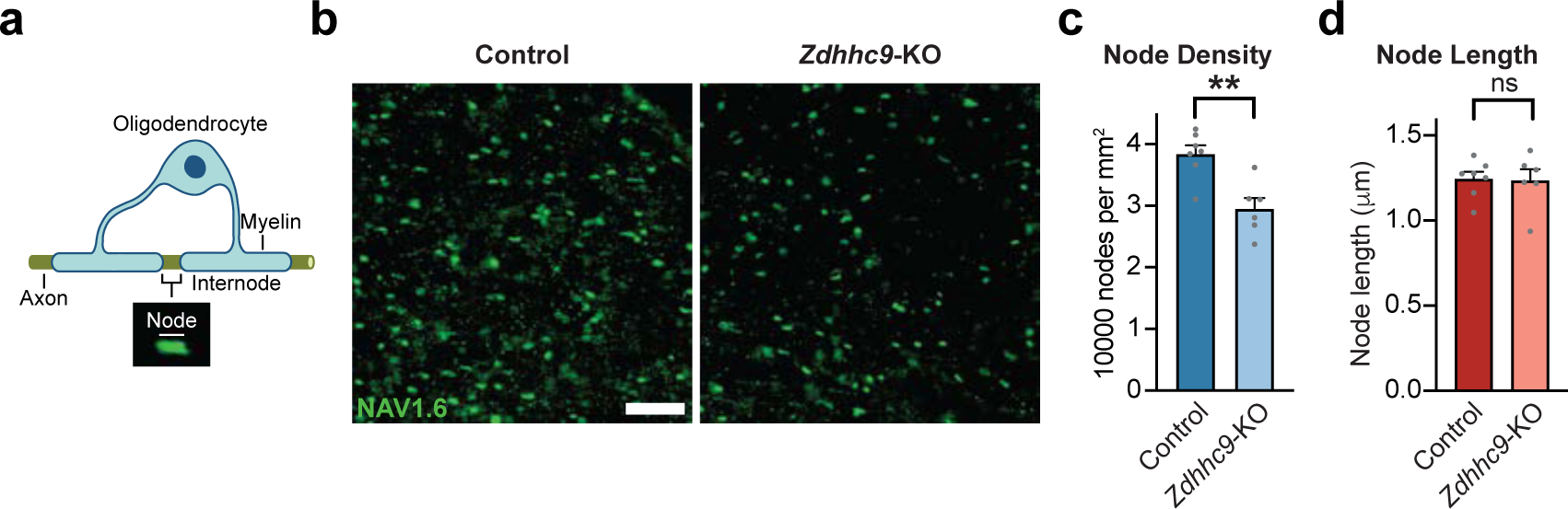
There is a reduction in the density of nodes of Ranvier in *Zdhhc9*-KO mice. **a** Illustration of a myelinated axon and node of Ranvier. **b** Representative images of NAV1.6 (nodal marker) immunostaining within the P60 corpus callosum (scale bar, 10μm). **c** Quantification of node density and **d** the node length (*n* = 6 *Zdhhc9*-KO mice, *n* = 7 control mice). Data in **c** and **d** represent mean + SEM, statistical tests were done using unpaired two-tailed Student’s t-tests.

**Fig. S7.**
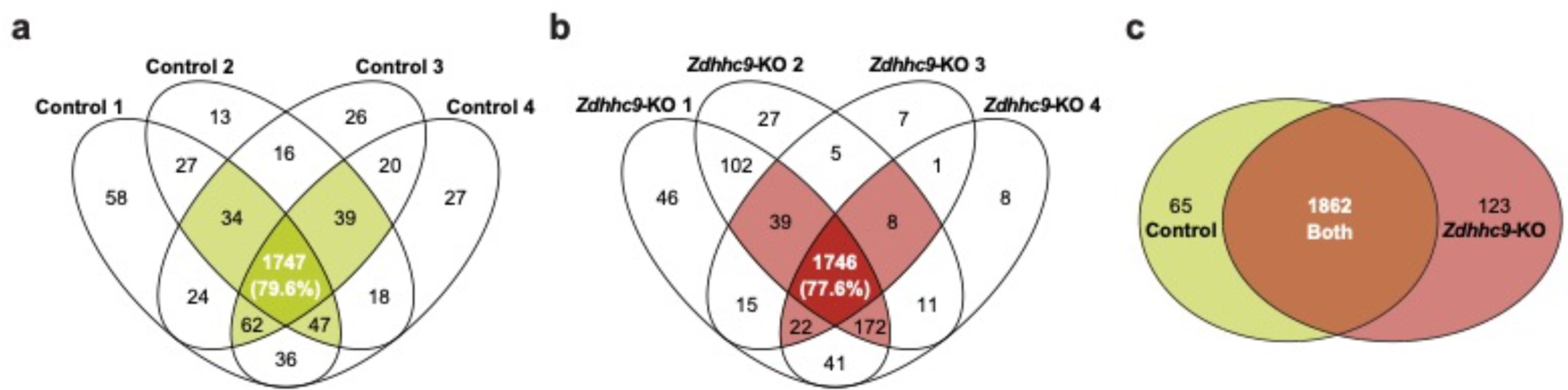
Analysis of proteins identified in both *Zdhhc9*-KO and control myelin samples. Venn diagrams showing the overlap of proteins identified in **a,** control and **b,** *Zdhhc9*-KO samples. **c** Venn diagram of myelin proteins identified in at least 3 *Zdhhc9*-KO and control samples.

